# A Nextflow pipeline for molecular quantitative trait loci mapping in small sample size datasets with an application in Atlantic salmon

**DOI:** 10.1101/2025.06.17.660077

**Authors:** Dat Thanh Nguyen, Simen R. Sandve, Sigbjørn Lien, Lars Grønvold

## Abstract

**Background:** Molecular quantitative trait loci (molQTL) mapping, particularly for gene expression and chromatin accessibility, provides crucial insights into the regulatory and functional potential of genetic variation. While significant progress has been made in humans and model organisms, aquatic genomics remains underexplored due to the large sample sizes typically required for statistical power.

**Results:** In this work, we enhance the scalability, reproducibility, and accessibility of the well-established RASQUAL method, which efficiently detects molQTLs in small datasets, by leveraging the Nextflow workflow framework. This adaptation, named nf-RASQUAL, supports fully automated QTL mapping and incorporates a robust, comprehensive multiple-testing correction process. We apply the pipeline to a comprehensive multi-omics dataset from Atlantic salmon, identifying numerous significant expression and chromatin accessibility QTLs across multiple tissues. Our analysis reveals that a large proportion of lead variants for these loci reside in non-coding regions, with caQTL lead SNPs more likely to disrupt transcription factor motifs. Additionally, the enriched colocalization of eQTL and caQTL lead SNPs in brain, liver, and gonad tissues suggests shared regulatory mechanisms.

**Conclusions:** These findings highlight the scalability and utility of nf-RASQUAL for advancing genetic regulation research in aquaculture, facilitating molQTL studies in less-explored species, and improving our understanding of molecular phenotypes shaped by genetic diversity.

## 1 Background

Genome-wide association studies (GWAS) have successfully identified myriad genetic variants linked to human diseases as well as complex agricultural and aquacultural traits[1, 2, 3]. However, the fact that associated loci are mostly situated in non-coding regions of the genome makes understanding the underlying gene regulation mechanisms nontrivial [4, 5]. Among various approaches, mapping genetic variants to molecular traits (molQLT) including cellular phenotypes such as gene expression (eQTL), splicing (sQTL), and chromatin accessibility (caQTL) is a particularly effective way to investigate the regulatory potential of GWAS-associated variants [6]. Indeed, numerous molQLT studies investigating various molecular traits have identified abundant QTLs associated with gene expression, [7, 8, 9, 10, 11, 12], alternative splicing [13], RNA editing [14, 15], circular RNAs [16, 17], DNA methylation [18, 19, 20], and chromatin accessibility [21, 22, 23, 24] that offer precise information on molecular functions influenced by human genetic variation.

As the human population expands, the need to generate an ample supply of nutritious food using fewer natural resources becomes urgent. This is essential not only to mitigate hunger and malnutrition but also to minimize the environmental footprint of animal farming and preserve biodiversity. To achieve this objective, comprehending the genetic control of molecular phenotypes in agricultural species, including farm animals holds the promise of enhancing production traits through genetics-based approaches [25, 26]. Despite the recent successes of molQLT in cattle and pigs [27, 26], the genetic regulation of molecular phenotypes in aquaculture species remains poorly characterized. To reduce the knowledge gap in aquatic species, the AQUA-FAANG consortium has recently generated a large number of multi-tissue matched genotypes and phenotype datasets for six major farmed fish species in Europe [28]. Despite its comprehensiveness, a primary obstacle to applying molQLT to the AQUA-FAANG project datasets is the requirement for large sample sizes in association testing, given the modest effect sizes of common variants [22].

To address this issue, we develop a scalable and reproducible pipeline named nf-RASQUAL for efficient molQLT mapping, leveraging Nextflow [29] and RASQUAL (Robust Allele-Specific Quantitation and Quality Control) [22]. RASQUAL is a probabilistic framework known for its effectiveness in association mapping of molecular phenotypes, particularly in datasets with small or modest sample sizes [22, 30, 31, 32]. By applying the new computational pipeline to a multi-tissue sequencing dataset of Atlantic salmon, which includes whole genome, RNA-seq and ATAC-seq datasets covering five tissues including brain, gonad, liver, muscle, and gill; we identify a substantial number of eQTLs and caQTLs.

## 2 Methods

### 2.1 Workflow and implementation

An overview of the nf-RASQUAL pipeline is presented in Figure 1. The pipeline firstly processes inputs including genotype data (VCF), read mapping data (BAM), metadata, and a read count matrix to prepare for QTL mapping. It then performs QTL mapping with RASQUAL and finally, a multiple testing correction is employed to handle false discovery rate (FDR).

**Figure 1.**
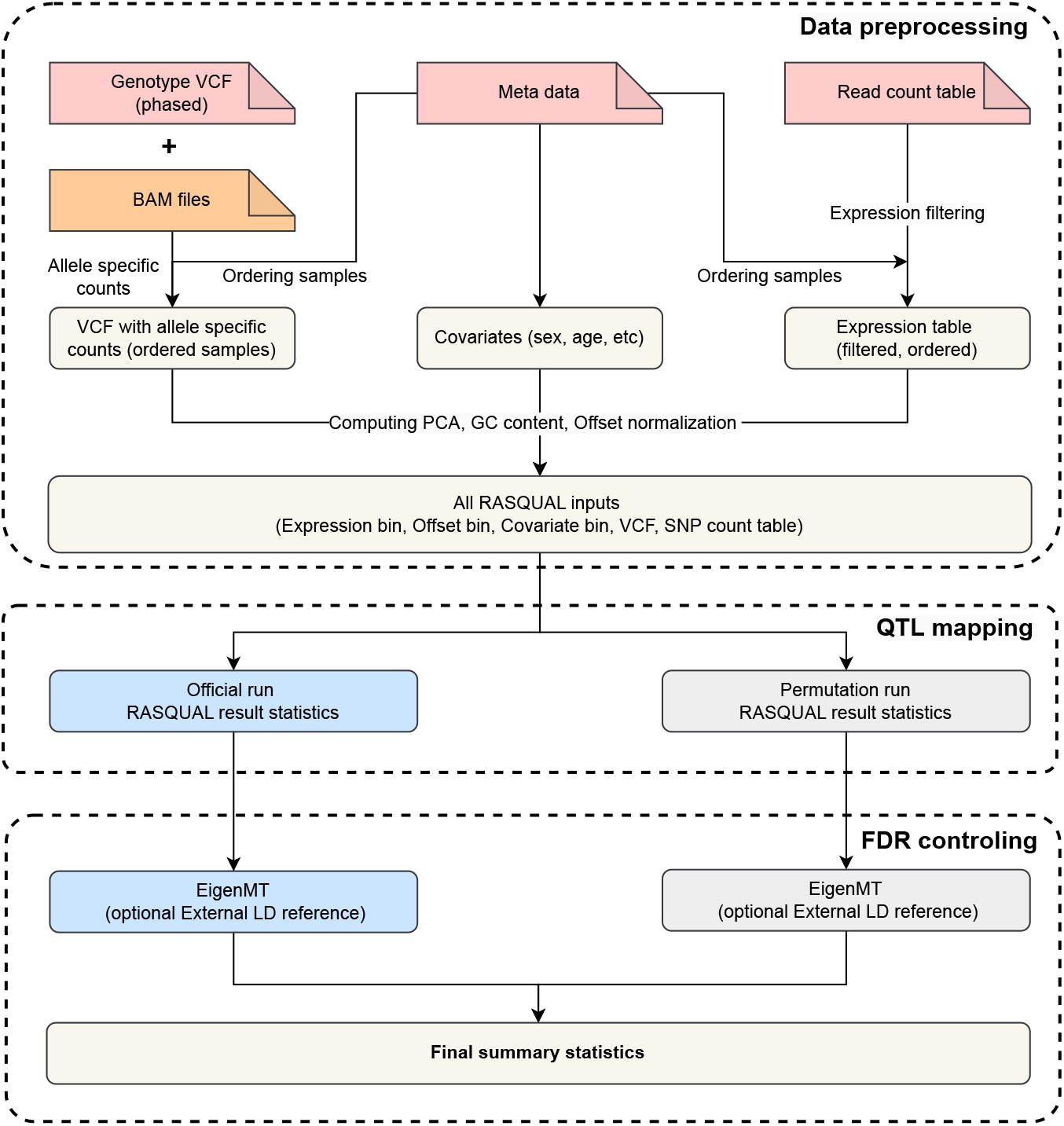
Overview of the nf-RASQUAL pipeline. The pipeline begins by integrating allele-specific information from matched BAM files, alongside processing metadata and expression tables to generate the input required for RASQUAL. QTL mapping is then performed, followed by multiple testing correction with EigenMT. An identical analysis for permuted data is employed to control false discovery.

In the first step, the genotype data is reordered to match the metadata and combined with BAM file information to generate allele-specific VCF files using the `createASVCF.sh` script from RASQUAL [22]. Simultaneously, the read counts for molecular phenotypes (RNA-seq and ATAC-seq) are normalized based on the recommended protocol of RASQUAL. We then perform a filter to retain features expressed in at least 50% of samples.

The phenotype data is then subjected to principal component analysis (PCA) to account for possible confounding factors. Due to the relatively small sample size in our dataset, we set the default option to only keep the two largest principal components and add these as covariates together with the metadata (such as sex and age, etc). To adjust for technical artifacts and feature-specific biases, the read count and GC content of the tested features are used to compute the offset using the `rasqualCalculateSampleOffsets` function from the `rasqualTools` package [24]. The expression, offset, and covariate data are converted into binary format compatible with RASQUAL using the `saveRasqualMatrices` using the same package. Next, the `countSnpsOverlapingExons` function is used to extract additional RASQUAL inputs, including feature ID, chromosome name, strand, feature start and end positions, testing window boundaries, the number of feature SNPs, and the total number of SNPs in the testing region. In this implementation, default testing windows are set to 10 kb for ATAC-seq and 500 kb for RNA-seq data.

After preparing all necessary input files, QTL testing is conducted in parallel using RASQUAL through a custom job scheduler. Since QTL testing involves multiple comparisons, we integrate EigenMT [33] into the pipeline to estimate the number of independent tests for each phenotype feature (e.g., genes or chromatin peaks) by leveraging population linkage disequilibrium (LD) data. Customized scripts are used to process input genotypes and RASQUAL outputs, making them compatible with EigenMT. Notably, using an external LD reference from a larger, genetically similar population is optional but recommended to improve the accuracy of independent test estimates. This recommendation arises because larger populations provide more robust LD estimates, and a genetically aligned reference population ensures the estimates remain relevant and reliable [33]. Bonferroni correction is then applied to compute adjusted p-values. Additionally, following the approach of Alasoo et al. (2018), one additional run of RASQUAL with the --random-permutation option is performed to generate empirical null p-values based on permuted sample labels. The same EigenMT procedure is then applied to these permuted p-values, allowing a comparison of the true association p-values with the empirical null distribution to identify QTLs with certain FDR cutoff.

### 2.2 Dataset and data preprocessing

To assess the effectiveness of the proposed workflow, we utilize a multi-omics dataset of Atlantic salmon provided by the AQUA-FAANG consortium [28]. This dataset includes genome sequencing, RNA-seq, and chromatin accessibility (ATAC) data from twelve fish, sampled across five key tissues: brain, gonad, liver, muscle, and gill (with no ATAC-seq data available for the gill). All data were uniformly preprocessed using nf-core pipelines [34]. In brief, quality control was applied to all sequencing data, followed by alignment to the Atlantic salmon reference genome v3.1 [35] using BWA [36] for whole-genome and ATAC-seq data, and STAR [37] for RNA-seq data. Variant calling was performed using HaplotypeCaller [38], transcript abundance was quantified using salmon software [39] from STAR-aligned [37] BAM files, and peak calling was conducted with MACS2 [40]. FeatureCounts [41] was used to quantify fragments overlapping consensus chromatin accessible peak (caPeak) annotations. Before conducting eQTL and caQTL analyses, the genotype data are further processed by phasing with Beagle5 [42]. We also apply a filter to retain SNPs with a minor allele frequency greater than 1% and Hardy-Weinberg equilibrium (HWE) p-values *>* 1e-6, based on a large-scale population data of wild Atlantic salmon [43].

### 2.3 Motif disruption analysis

We explored the characteristics of caQTL lead variants by conducting motif disruption analyses across the four tissues examined. To do this, we extracted caQTL lead variant information including genomic positions, reference allele and alternative allele for motif disruption tests. In cases of ties, we randomly selected one variant. Next, we used the `pysam` package to retrieve 30 bp DNA sequences centered on each SNP position for both reference and alternative alleles.

These sequences are then scanned against transcription factor (TF) motifs from the JASPAR 2020 CORE vertebrates non-redundant database [44] using Find Individual Motif Occurrences (FIMO) implemented in the MEME Suite v5.5.7 [45, 46]. Following a protocol proposed by Currin et al. [32], we apply FIMO with parameters `--thresh 0.01; --max-stored-scores 1000000; --no-qvalue; --skip-matched-sequence;` and `--text`. We only keep motif occurrences that overlapped caQTL variant positions. We further apply filtering to retain motif-variant pairs with at least one allele passing the significant cutoff of 1e-4 which is the recommended cutoff of FIMO for motif identification [45].

## 3 Results

### 3.1 Discovery of eQTLs and caQTLs in Atlantic salmon

Using nf-RASQUAL, we conduct eQTL association tests for approximately 1.4 million SNPs against 26,708; 25,666; 19,406; 18,629; and 25,960 expressed genes for in the brain, gonad, liver, muscle, and gill, respectively. Regarding caQTL, the number of caPeaks tested were 175,099; 317,837; 240,388; and 324,642 for the brain, gonad, liver, and muscle. In parallel with the experimental data, an identical procedures are applied to the corresponding permutation data to assess the robustness of the method and control for false discovery.

To validate the effectiveness of the proposed computational pipeline in identifying associations between genetic variants and molecular phenotypes, we extract the lead SNPs (smallest p-values) for all tested features from both the observed and permuted datasets. The p-value pairs are visualized as QQ plots, illustrating the relationship between -log_10_(*p*) values for observed versus permuted data as shown in Figure 2 A and 2 B. In both plots, the genome-wide distribution of test statistics for the observed data is substantially higher than the expected permuted distribution, especially in the right tail, for all tissues analyzed. This indicates no evidence for systematic spurious associations in the observed data, confirming the reliability of the results.

**Figure 2.**
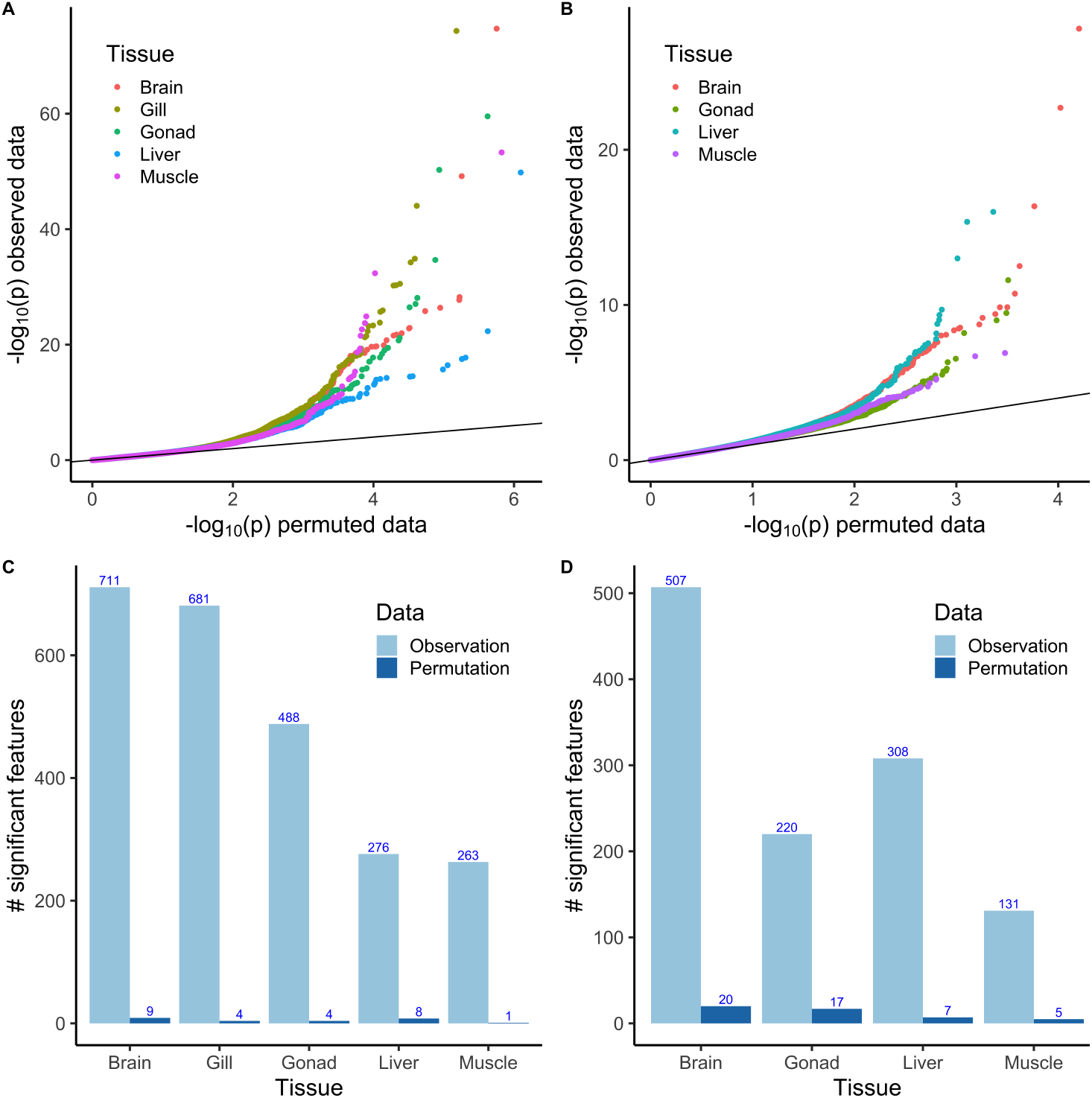
eQTL and caQTL discovery in Atlantic salmon. A, B Q-Q plot of lead p-values in observed versus permuted data of eQTL and caQTL respectively, the black solid line is the diagonal. C, D Bar plot comparing the number of significant features identified in observed versus permuted data for eQTL (C), and caQTL (D).

To account for multiple testing, we utilize the obtained population data of wild Atlantic salmon [43] as a reference panel for independent tests estimation by EigenMT [33]. The Bonferroni correction procedure is then applied with a cutoff of 10%. Figure 2 C and 2 D present bar charts showing the final number of significant genes (eGenes) or caPeaks (ePeaks) identified in both the observed and permuted data across different tissues. Figure 2 C specifically highlights the number of eGenes identified in the eQTL analysis. In the observed data, the brain exhibits the highest number of significant features (711), followed by the gill (681), while the gonad (488), liver (276), and muscle (263) display fewer number of eGenes. In contrast, the permuted data show only a small number of significant features across all tissues, with the brain having the highest count (9), indicating that significant signals in the observed data are not artifacts of random variation.

Figure 2 D displays the caQTL results, comparing the number of ePeaks in the observed and permuted data. In the observed data, the brain again shows the highest number of ePeaks (507), followed by the gonad (220), liver (308), and muscle (131). Like with the eQTLs, the permuted data have only a very small number of significant caPeak, suggesting that the significant caQTLs are likely driven by biological relevance rather than random noise.

### 3.2 Characteristics of molQLT variants in Atlantic salmon

To investigate molQLT variants, we predict the effect of lead variants (selecting one randomly in cases of ties) from all eGenes and ePeaks using the Ensembl Variant Effect Predictor (VEP) [47].

As shown in Figure 3, the six most abundant categories of genetic variants across tissues and QTL types are located in introns, intergenic regions, upstream and downstream of genes, and within the 3’ and 5’ untranslated regions (UTRs). Intron variants consistently dominate across all tissues, representing the majority of identified variants. For instance, they account for 59.76% of brain caQTLs, 58.51% of brain eQTLs, 57.42% of gill eQTLs, 62.27% of gonad caQTLs, and 61.48% of gonad eQTLs lead SNPs. Liver tissues show slightly lower proportions of intron variants, with 54.55% in caQTLs and 53.99% in eQTLs while intron variants comprising 49.62% of and 58.56% of muscle caQTL and eQTLs lead SNPs respectively.

**Figure 3.**
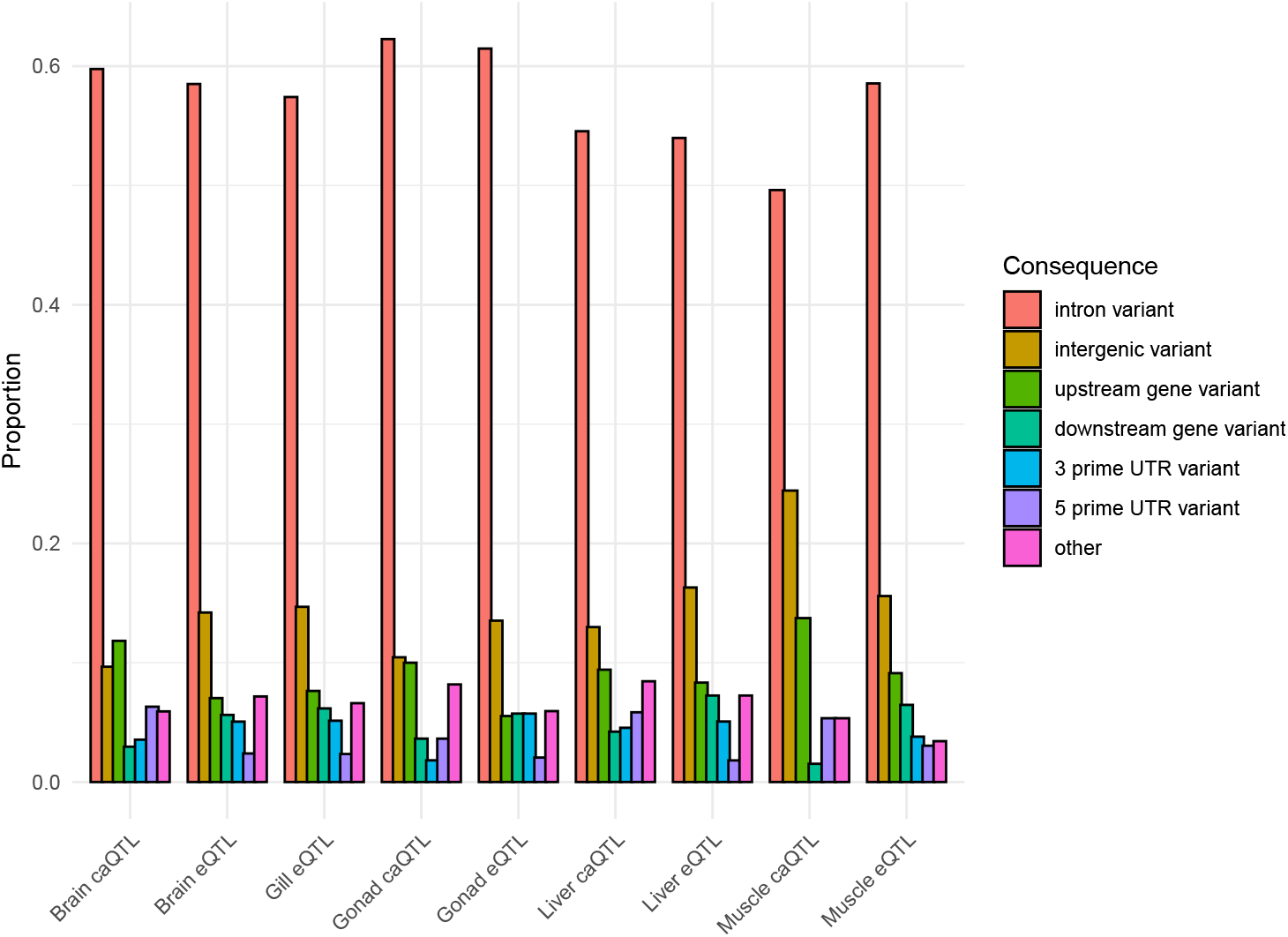
Ensembl Variant Effect Predictor annotation of molQLT variants across tissues (brain, gill, gonad, liver, and muscle) and QTL types (caQTLs and eQTLs).

Intergenic variants emerge as the second most common category but display variability across tissues and QTL types. In brain tissue, intergenic variants constitute 9.66% of caQTLs and 14.21% of eQTLs, while in gill eQTLs they account for 14.68% of lead SNPs. Gonad tissues show slightly lower proportions, with intergenic variants representing 10.45% of caQTLs and 13.52% of eQTLs lead SNPs while muscle caQTLs stand out with a notably higher proportion of intergenic variants at 24.43%. Upstream gene variants are less frequent but show consistent representation across tissues, with proportions ranging from approximately 5–13% of lead SNPs. For example, they account for 11.83% of brain caQTLs and 7.03% of brain eQTLs. Variants in downstream regions and untranslated regions (both 3’ and 5’ UTRs) occur less frequently, typically comprising 2-7% of all variants across tissues and QTL types.

Overall, these results demonstrate that intron variants are the predominant type across all tissues and QTL types, suggesting their central role in regulatory processes. However, the variability observed in intergenic and other variant types indicates tissue-specific differences in genetic regulatory architectures.

### 3.3 Motif disruption analysis

We measured the difference in motif matching between variant alleles by calculating the log ratio of FIMO p-values followed by previous studies [48, 32]. The FIMO p-values are defined as the probability of a random sequence of the same length as the motif matching that position of the sequence with an equal or higher score [45]. For each variant-motif pair, motif disruption is calculated as *log*10(*paw*) – *log*10(*pas*), where *paw* and *pas* are the FIMO matching p-values for weaker and stronger alleles. A motif is defined as disrupted if the disruption value exceeds 1, indicating a 10-fold difference in FIMO p-values between alleles.

We analyze the motif disruption rates of lead SNPs between ePeaks and non-ePeaks across 746 non-redundant JASPAR motifs in brain, gonad, liver, and muscle tissues. For ePeaks, we test 506, 220, 306, and 131 variants and observe 562, 217, 363, and 163 disruptive events in brain, gonad, liver, and muscle tissues, respectively. To evaluate the statistical significance of these findings, we conduct a ratio z-test comparing the disruption rates of ePeaks to those of non-significant caPeaks.

The results indicate significant enrichment in motif disruption rates for brain (p = 0.010), liver (p = 0.001), and muscle (p = 0.006) tissues when comparing significant peaks to the baseline disruption rates of non-significant peaks. In contrast, no statistically significant difference is observed in gonad tissue (p = 0.890). To facilitate interpretation and visualization of these results, we normalize the number of disruptive events by the number of variants tested for each tissue type (Figure 4). This normalization reveals a moderate enrichment in motif disruption rates for ePeaks relative to non-ePeaks in brain (1.11 vs. 0.996), liver (1.19 vs. 1.004), and muscle (1.24 vs. 1.004). However, gonad tissue does not show such enrichment (0.986 vs. 0.996). These results suggest that caQTL lead SNPs have a higher likelihood of disrupting TF motifs compared to non-significant SNPs in brain, liver, and muscle tissues but not in gonad tissue.

**Figure 4.**
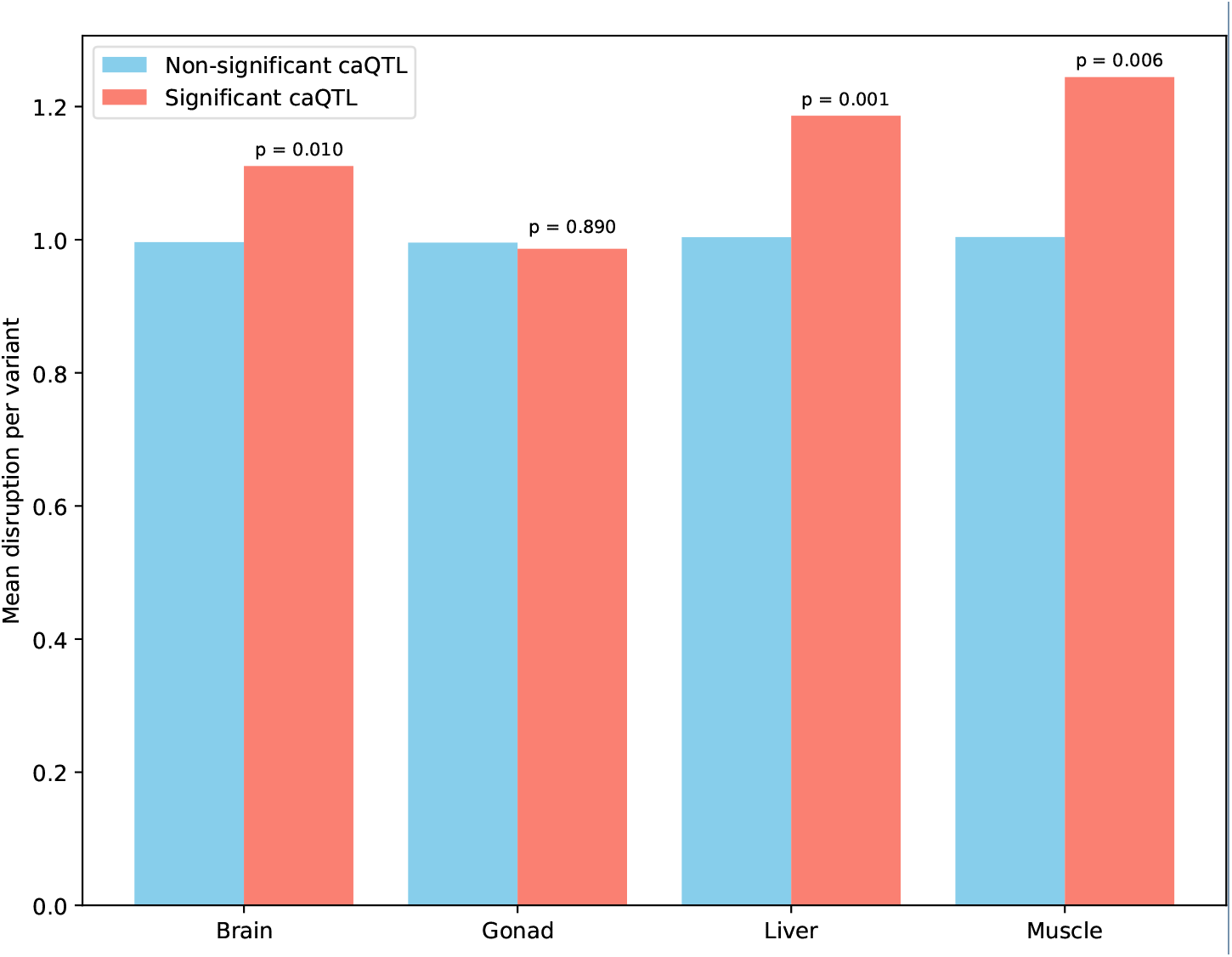
Normalized motif disruption rates for significant and all caPeaks across tissues.

### 3.4 caQTL are enriched in eQTL loci

We investigate the colocalization between eQTL and caQTL across brain, gonad, liver, and muscle tissues. Specifically, we calculate LD between eGenes and ePeaks lead SNPs using PLINK v1.9 [49], leveraging previously described population data [43]. Colocalization is defined when lead SNPs have strong pairwise LD (*r*^2^ *>* 0.7). The baseline colocalization rate is determined as the ratio of colocalized loci to all tested caPeaks. This rate is then compared with the colocalization ratio of significant caPeaks, defined as the ratio of colocalized loci to all significant caPeaks. We perform permutation tests under the null hypothesis that the baseline colocalization rate represents the expected colocalization rate.

Table 1 presents the colocalization rates and enrichment ratios in various tissues. We observe strong enrichment in brain, gonad, and liver tissues but not in muscle. The baseline colocalization rates range from 0.07% in muscle to 0.18% in brain, while the colocalization rates for significant caQTLs are notably higher in brain (1.18%), gonad (1.36%), and liver (0.97%). Statistical significance (permutation test *p <* 0.05) is observed in these three tissues, with substantial enrichment ratios of 6.42, 10.20, and 10.23 respectively, indicating enhanced colocalization beyond baseline expectations. In contrast, muscle tissue shows no enrichment, with zero colocalization events detected among 131 significant caQTLs, likely due to the limited number of ePeaks identified.

**Table 1.**
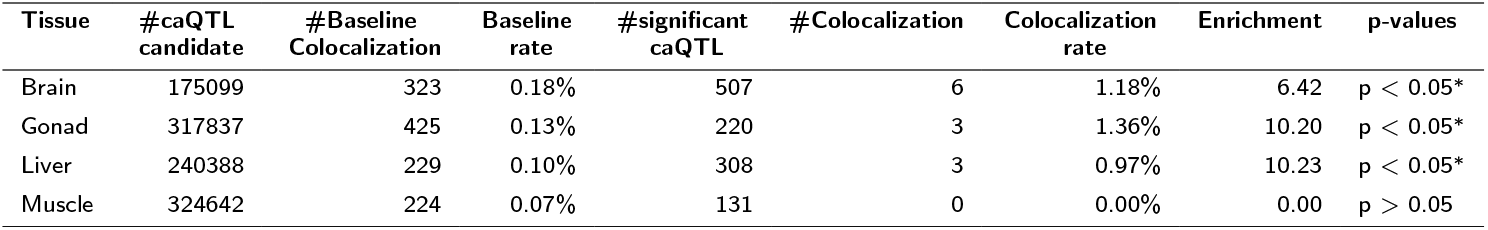
Colocalization rates and enrichment in various tissues. ^*∗*^ indicates statistical significant in a permutation test under the null hypothesis is the baseline colocalization rate

| Tissue | #caQTL candidate | #Baseline Colocalization | Baseline rate | #significant caQTL | #Colocalization | Colocalization rate | Enrichment | p-values |
| --- | --- | --- | --- | --- | --- | --- | --- | --- |
| Brain | 175099 | 323 | 0.18% | 507 | 6 | 1.18% | 6.42 | p < 0.05* |
| Gonad | 317837 | 425 | 0.13% | 220 | 3 | 1.36% | 10.20 | p < 0.05* |
| Liver | 240388 | 229 | 0.10% | 308 | 3 | 0.97% | 10.23 | p < 0.05* |
| Muscle | 324642 | 224 | 0.07% | 131 | 0 | 0.00% | 0.00 | p > 0.05 |

## 4. Discussion

QTL mapping of genetic variants with intermediate molecular phenotypes has been established as a robust approach for understanding regulatory mechanisms of genetic variants. Despite its potential, a major challenge in applying QTL mapping lies in the need for large sample sizes due to the typically modest effect sizes of common variants. This limitation is particularly pronounced in studies with smaller sample sizes, reducing the power to detect significant associations. In this study, we extend the scalability, reproducibility, and user-friendliness of the well-established RASQUAL method [22] by leveraging the Nextflow workflow framework [29]. The developed pipeline further enables efficient QTL mapping in a fully automated manner, with an integrated comprehensive multiple-testing correction procedure using EigenMT [33].

To illustrate the usability and effectiveness of the developed pipeline, we apply it to a multi-omics dataset of Atlantic salmon spanning five key tissues. Through rigorous comparison with permutation tests, we demonstrate that the pipeline successfully identifies hundreds of significant eQTLs and caQTLs across tissues. Variant annotation reveals that lead molQLT variants in Atlantic salmon predominantly reside in non-coding regions, with a high proportion in intron and intergenic regions, consistent with findings from molQLT studies in other species [11, 27, 26]. Furthermore, motif disruption analysis suggests that lead variants associated with significant caPeaks are more likely to disrupt transcription factor motifs in brain, liver, and muscle tissues compared to non-significant caPeaks, highlighting the functional impact of caQTL variants.

The colocalization analysis of eQTL and caQTL lead SNPs reveals enriched colocalization in brain, gonad, and liver tissues, suggesting shared regulatory mechanisms influencing both gene expression and chromatin accessibility in these tissues. This colocalization points to potential causal variants driving both eQTL and caQTL signals. Although colocalization in muscle tissue does not reach statistical significance, likely due to limited statistical power, the observed enrichment in other tissues supports the biological relevance of these findings.

## 5 Conclusions

Overall, this study introduces nf-RASQUAL and demonstrates its reliability and scalability for QTL mapping across multiple tissues. The integration of motif disruption and colocalization analyses enhances result interpretation, offering deeper insights into the regulatory mechanisms shaped by genetic variation in Atlantic salmon. With its fully automated workflow and capacity for large-scale testing, we believe the nf-RASQUAL pipeline will be a valuable tool for investigating genetic regulation across various biological systems, particularly in aquatic species studied under the AQUA-FAANG project.

## Availability of data and materials

The nf-RASQUAL pipeline is freely available at https://github.com/datngu/nf-rasqual. Data supporting the findings and source codes for data analyses and generating figures for this study are available at https://github.com/datngu/nf-RASQUAL_paper.

## Use of AI Software

Large language models were used to improve the wording and grammar of some texts, but not to generate new content.

## Acknowledgements

The authors would like to thank the AQUA-FAANG consortium for granting access to the dataset. We also extend our gratitude to the Orion Cluster for providing computational resources and to Teshome Dagne Mulugeta for technical support with server-related issues.

## Abbreviations

SNP: Single Nucleotide Polymorphism
GWAS: Genome Wide Association Studies
eQTL: expression quantitative trait loci
caQTL: chromatin accessibility quantitative trait loci
MAF: Minor Allele Frequency

## Ethics approval and consent to participate

Not applicable.

## Consent for publication

Not applicable.

## Author details

Centre for Integrative Genetics, Faculty of Biosciences, Norwegian University of Life Sciences,, Norway, 1432 Ås, Norway.

